# Integrative signatures of signaling pathway response increase robustness and accuracy of pathway predictions

**DOI:** 10.1101/2022.06.03.494712

**Authors:** Nicholas A. Clark, Yan Ren, David R. Plas, Siva Sivaganesan, Mario Medvedovic

## Abstract

**Motivation:** Aberrant cell signaling is known to drive progression of cancer and many other diseases. The study of signaling pathways within cells is central to identifying drugs that seek to modulate these pathways. Expression of pathway genes (i.e. genes that code for pathway proteins) correlates poorly with signaling pathway activity, making prediction of signaling pathway activity changes based on transcriptional disease signatures a challenging problem. Pathway architecture and response also varies across cell lines, which reflects how drug response varies across a patient population.

**Results:** Here, we present a transcriptional footprinting framework for predicting changes in activity of signaling pathway by integrating transcriptional signatures of genetic perturbations of pathway genes over a diverse set of cell lines into a integrative Pathway Activity Signature (iPAS). We use an unsupervised multi-task learning approach to create pathway signatures across 12 cell lines using genetic loss of function data from the LINCS project. We also use supervised learning to construct an optimal predictor based on the ensemble of 12 cell line signatures. Our methods achieve a sizeable increase in performance, as measured by prediction of pathways targeted by LINCS chemical perturbagens.

**Availability:** Open source R package iPAS is available at https://github.com/uc-bd2k/iPAS.

**Supplementary information:** Supplementary data are available online.

## Introduction

Signaling pathway alterations drive pathobiology in cancer and other diseases (Hanahan and Weinberg, 2000; Finkel and Gutkind, 2003; Saxton and Sabatini, 2017). Mechanistically, a signaling pathway consists of a network or cascade of interacting proteins. These proteins can be covalently modified in a number of ways by other intra-cellular proteins, which can alter their physical conformation, stability, and/or enzymatic activity. The end result of activation or inhibition of a signaling pathway is often an alteration in the transcriptional state of the cell.

It is possible to directly measure signaling protein and pathway activity using proteomic technologies such as RPPA and mass-spectrometry. Nevertheless, the low-cost and convenience of transcriptomic profiling led to it becoming the dominant platform for genome-wide functional analysis. Large-scale databases of transcriptomic response to genetic and chemical perturbagens, such as those produced by the LINCS project (Subramanian et al., 2017; Pilarczyk et al., 2019), offer an extraordinary resource from which to learn genes that are predictive of pathway response.

Standard methods for pathway activity analysis using transcriptomic data rely on assessing the changes in gene expression of genes coding for pathway proteins, derived from sources like Gene Ontology (GO) (Gene Ontology Consortium, 2017), Reactome (Jassal et al., 2019), and KEGG (Kanehisa et al., 2017). However, changes in signaling pathway activity correlate poorly with changes in expression of pathway genes (Geistlinger et al., 2016; Dugourd and Saez-Rodriguez, 2019). To overcome this obstacle, recent methods such as SPEED (Parikh et al., 2010; Rydenfelt et al., 2020), PROGENy (Schubert et al., 2018), the MSigDB Hallmark collection (Liberzon et al., 2015), and pasLINCS (Ren et al., 2020) have used transcriptional “footprints” (Dugourd and Saez-Rodriguez, 2019), the changes in genes that *respond* to pathway manipulation, to predict pathway activity.

Tumor-derived cell lines serve as model systems for cancers originating from specific anatomical and histological contexts (Goodspeed et al., 2016; Mirabelli et al., 2019). Driver mutations may be specific to each individual tumor and cell line but activated pathways responsible for cancer hallmarks are often shared or overlapping. Moreover, the proteins involved in these pathways and the architecture of their interaction under normal conditions are largely similar in most cells. Here, we present an approach that integrates the transcriptional responses and pathway topology across an array of cell lines to create what we call integrative Pathway Activity Signatures (iPAS). iPAS signatures are then used to detect changes in activity of signaling pathways in disease and other transcriptomic signatures.

We benchmarked the performance of our iPAS signatures by measuring how well we could predict the KEGG pathway targeted by small molecule inhibitors in the LINCS dataset. In previous benchmarks, the method integrating pathway topology and transcriptional signatures of genetic perturbation of the genes in the pathway in a single cancer cell line outperformed all competing methods tested (Ren et al., 2020), including the topological pathway analysis methods such as CePa (Gu and Wang, 2013), SPIA (Tarca et al., 2009), and PathNet (Dutta et al., 2012), as well as simple enrichment approaches. iPAS integrates data from 12 different cell lines using multi-task and ensemble learning. The multi-task method seeks to exploit similarity in pathway response across cell lines, while the ensemble method seeks to exploit heterogeneity in response. We find both methods increase the accuracy of our predictions in comparisons to using a single cell line at a time.

## Methods

### Methodology overview

iPAS methodology identifies pathway response genes that are predictive of altered signaling activity by genetic or chemical perturbation. It creates cell line-specific pathway signatures and a framework for making consensus predictions of altered pathway activity across different cell lines. We create pathway signatures using LINCS genetic loss of function perturbation (GP) signatures of of pathway proteins along with pathway topology information from KEGG. We incorporate GPs from 12 cell lines in the signature creation process. Figure 1 shows an overview of the methodology.

**Figure 1:**
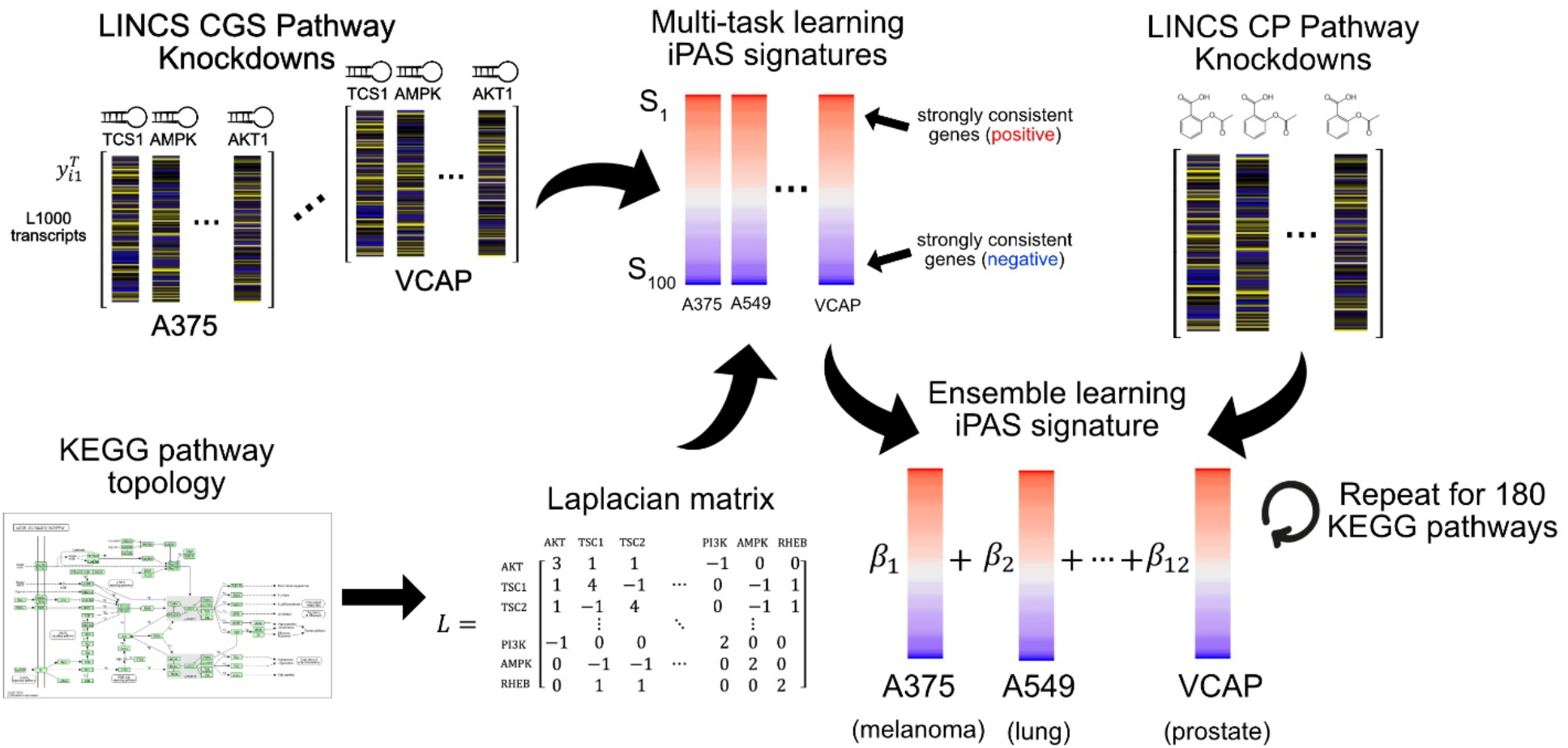
Overview of ensemble signature creation for iPAS. Individual iPAS signatures are first created for each cell line, using a “pathway consistency score” based on the signed Laplacian representing pathway topology. All pathway signatures (for each pathway, for 12 cell lines) are then correlated with CPs of drugs targeting one pathway. These correlations are then used in a machine learning algorithm (logistic regression with a LASSO penalty) to solve for optimal combinations of the signatures from each cell line. This process of training with LINCS CP signatures is repeated for each of 181 pathways to create “iPAS ensemble” signatures for each pathway.

The framework incorporates two methods for data integration. Multi-task learning (Caruana, 1997; Yuan et al., 2016) balances the cell line-specific pathway response in 12 cell lines with average pathway response over those cell lines. The linear ensemble supervised learning (Wolpert, 1992; Naimi and Balzer, 2018) is used to predict affected pathways in a disease or a chemical perturbation signature based on similarity with the optimal consensus accross all cell lines. The ensemble predictor is trained using signatures of drugs that target the pathway from the LINCS chemical perturbagens (LINCS CP) dataset.

### Multi-task learning signatures of pathway activity

iPAS is based on the minimization of the multi-task learning functional where the regularization term favoring solutions that are consistent with the pathway topology and the muti-task penalty term favoring similar solution vectors for all cell lines. For a given pathway with GP signatures of *P* pathway nodes, we consider the expression of each gene separately. We let *y*_*ik*_ be the vector of expression of gene *i* in the *k*-th cell line across all *P* pathway nodes, 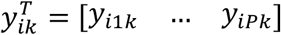 We look at its expression across all *N* cell lines: *y*_*i*_ = *vec*(*Y*_*i*_) where *Y*_*i*_ = [*y*_*i*1_ … *y*_*iN*_]. *Y*_*i*_ is a matrix where each column is the expression of gene *i* across pathway nodes in one cell line and *y*_*i*_ is the “vectorized” version of this matrix, a vector in which the columns of *Y*_*i*_ are stacked on top of each other.

The common mean for all target vector is defined as 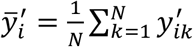. The “regularized” expression of gene *i*, in cell lin *k,ŷ*_*ik*_, is calculated by minimizing the corresponding regularization risk functional as follows:

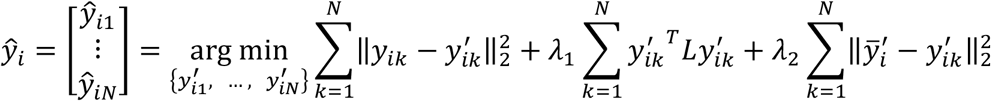

Where 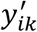 is a target vector in cell line *k* and {*ŷ*_*i*1_,…,*ŷ*_*iN*_} is the solution that minimizes the functional on the right-hand side.

The first term is the loss and keeps the target vector near the observed values, the second term if the regularization term favoring a vector that is consistent with the pathway topology, and the third term is the penalty that favors solutions that are close to the overall average across different cell lines. After jointly solving for *ŷ*_*ik*_ for each cell line for a range of regularization weights *λ*_1_, *λ*_2_ ∈ {0, 1, 2, 5, 10, 50, 100}, we define a pathway consistency score *S*_*ik*_ for each gene in each cell line based on its consistency with the pathway topology as defined by the Laplacian. In general, our pathway signature for each cell line 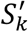 can be defined as a function of the vector of consistency scores for each gene (*S*_*k*_) and the regularized expression matrix of all genes (*Ŷ*_*k*_ = [*ŷ*_1_,_*k*_ … *ŷ*_978,*k*_]^*T*^): 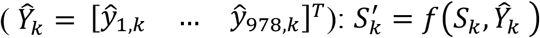 One reasonable option is to filter *ŷ*_*k*_ to the top *n* genes by score and then summarize this matrix into a vector, for example its first principal component. Another option is to simply use the consistency scores themselves as a signature of pathway activity.

The consistency score for gene *i* in cell line *k* is defined as the dot product with the first eigenvector of the pathway Laplacian 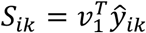 (Ren et al., 2020). For this score, vectors that are more similar to *v*_1_ will produce larger scores, which relate to higher consistency with the pathway. Low pathway consistency will result in scores near zero. Both negative and positive scores are possible, indicating that a gene is involved in response to the pathway, but in opposite directions, i.e. “activating” or “inhibiting” response. Each gene in each cell line is scored by its pathway consistency and 100 genes with the highest scores by absolute value are selected to serve as signature genes. The consistency scores for signature genes constitute the iPAS signature of the pathway activity.

Since pathway consistency score is the projection of *ŷ* on the first eigenvalue of the signed Laplacian L, known to belong to the null space of L, the choice of *λ*_1_ does not affect the values in our signature. Therefore, we only consider various values of *λ*_2_,the parameter that controls how much we push expression in each cell line toward average expression. When *λ*_2_ = 0,there is no sharing of information across cell lines and the signatures are essentially created separately. When *λ*_2_ = 100, the expression vector for each cell line is pushed almost completely to the mean over all cell lines.

### Cell-line agnostic consensus signatures of pathway activity

Pearson’s correlation is used as a measure of similarity between a query signature and a PAS signature for a pathway. Let *s*_*ijk*_ represent the Pearson’s correlation between query signature *i* (from *i* = 1, …, *N*) and the PAS for pathway *j* (from *j* = 1, …, *P*), in cell line *k* (from *k* = 1, …, *m* = 12). For each input signature and pathway, there are *m* = 12 measures of similarity (correlation). A linear predictor is used to combine the ensemble of similarity measures into *ŝ*_*ij*_ = *β*^*T*^*s*_*ij*_ = *β*_1_*s*_*ij*1_ + … + *β*_*m*_*s*_*ijm*_. Here, β is a vector of coefficients weighting the similarity to the PAS in each cell line.

The coefficients in the model were trained for each pathway separately using randomly selected two-thirds of the LINCS CP signatures targeting the pathway, and predictive ability is tested on the remaining third of the signatures. For pathway *j* we select all LINCS CP signatures that target the pathway. We then correlate these signatures with the PAS signatures for all pathways in all cell lines, giving us *m*-dimensional correlation vectors, or “points”, for each LINCS CP signature and each PAS. We consider the correlations with the PAS for pathway *j* as the “case” or “condition positive” group (giving a response of *y*_*i*_ = 0), while correlations with other pathway PASes are considered the “control” or “condition negative” group (giving a response of *y*_*i*_ = 1).

We construct a logistic regression problem using these two groups of points. This allows us to solve for a set of weights that best “separate” the two groups of points, giving high general similarity values *ŝ*_*ij*_ = *β*^*T*^*s*_*ij*_ = *β*_1_*s*_*ij*1_ + … + *β*_*m*_*s*_*ijm*_. for points in the “case” group and low general similarity values for points in the “control group”. The model is regularized using the LASSO penalty to encourage sparsity where possible, i.e. to drop cell lines where the PAS performs poorly.

### Benchmarking methods and ROC curves

iPAS was benchmarked by testing its ability to recognize signatures of drugs (LINCS chemical perturbagens -CP) that target the pathway. For multi-task iPAS signatures we use all CPs targeting the pathway and for ensemble signatures we use the portion of the pathway-targeting CP signatures (33%) that were held out from training as an independent test set. In each case, we use receiver operating characteristic (ROC) curves to summarize the performance of the signatures. ROC curves plot a line with sensitivity (also known as recall or true positive rate) on the y-axis and 1 – specificity (also known as false positive rate) on the x-axis. Sensitivity is the proportion of signatures “positive” for a condition (here, signatures that target a given pathway) that are correctly predicted as “positive”. Specificity is the proportion of signatures “negative” for a condition (here, signatures that do not target a given pathway) that are correctly predicted as “negative”.

The ROC curve is plotted for all possible prediction thresholds as the change in this threshold causes the false positive rate (on the x-axis) to range from 0 to 1. The area under the ROC curve (AUROC) summarizes the performance of the classifier over all possible classification thresholds. A classifier that is no better than random chance will give an AUROC of around 0.5 while a good classifier will give a value closer to 1.

The benchmarking procedure is as follows: 1) For pathway, calculate Pearson correlation of the iPAS signature with all pathway-targeting CPs. 2) Calculate Pearson correlation of iPAS for all other pathways with the same CPs. 3) The correlation values from step 1 are considered to be “positive” or “case” group. The correlation values from step 2 are considered to be “negative” or “control” group. 4) Ignore any values from the negative group where the node targeted by the CP also belongs to the pathway. 5) From the “case” and “control” (or “positive” and “negative”) groups of correlations, ROC curves are constructed and summarized by their AUROC. 6) This process is repeated for each KEGG pathway.

## Results

### LINCS CP benchmarking results (multi-task signatures)

We tested the ability of our iPAS methodology to identify the signaling pathway targeted by LINCS chemical perturbagen (CP) signatures. The Pearson correlation of a CP with a pathway signature was used as the evidence of perturbation of the pathway. For each KEGG pathway, we constructed ROC curves to determine the ability of these correlations to correctly identify the pathway targeted by CPs. Figure 2 shows a summary of the result for the multi-task iPAS signatures.

**Figure 2:**
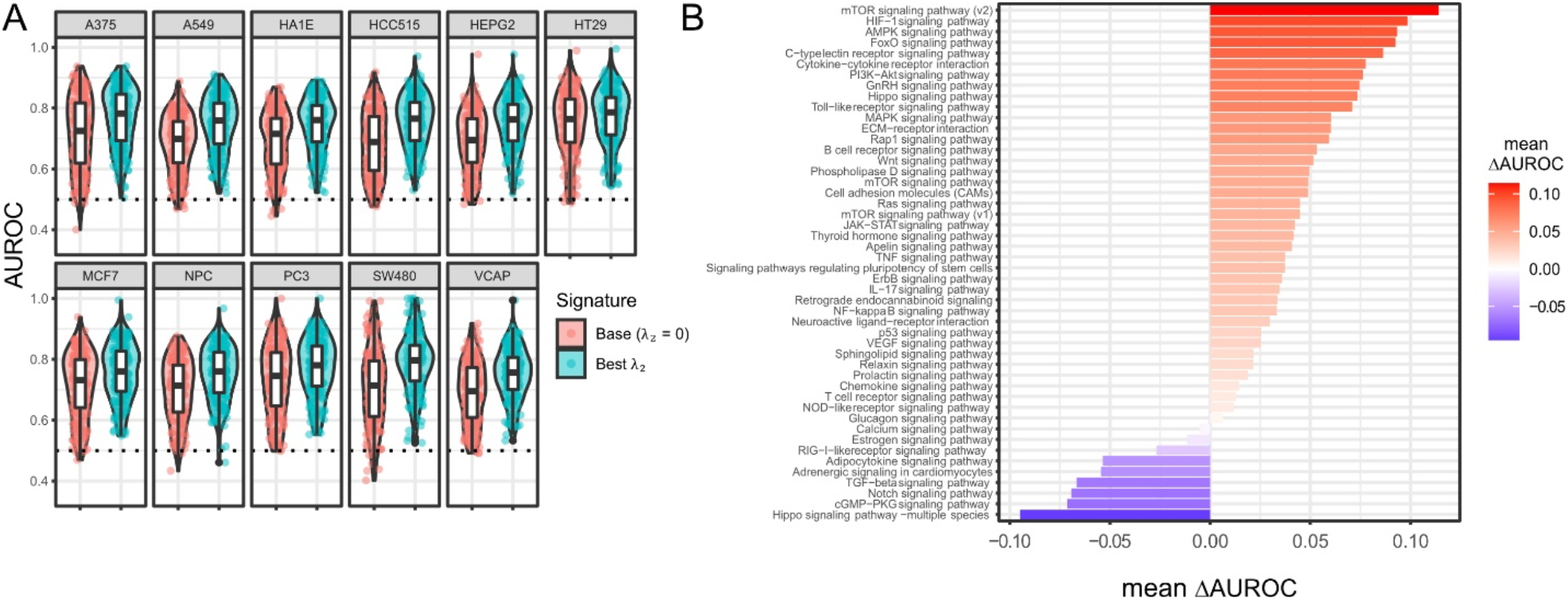
LINCS CP Benchmarking results for multi-task iPAS signatures. A) Box and violin plots of area under the curves (AUROC) for predicting which pathway is targeted by the chemical perturbagen. For each pathway we use iPAS signatures with the best performing *λ*_2_ value out of *λ*_2_ ∈ {1, 2, 5, 10, 50, 100}. We compare AUROC of iPAS signatures to signatures created separately in each cell line (*λ*_2_ = 0). B) Mean difference in AUROC across cell lines for iPAS signatures of signaling pathways from the lowest multi-task weight (*λ*_2_ = 0) to the highest multi-task weight *λ*_2_ = 100) (Not quite understanding this).

The box and violin plots in Figure 2A show the area under the ROC curve (AUROC) for all 181 pathways for signatures pertaining to each cell line, with red indicating *λ*_2_ = 0 (no sharing between cell lines) and blue indicating the AUROC of the best performing *λ*_2_ value. A larger area under the curve implies better prediction and an AUROC of 0.5 implies prediction that is no better than random chance. In each cell line, we see a statistically significant increase in AUROC from the base signatures (*λ*_2_ = 0) to the multi-task signatures (*λ*_2_ = *best value*). The full results for all *λ*_2_ values are shown in Fig. S3. We also looked at the difference between signatures made in each cell line with no data sharing (*λ*_2_ = 0) versus high data sharing (*λ*_2_ = 100) (Fig. 2B). We saw an increase in AUROC in most signaling pathways, with the highest increase in our curated mTOR pathway (v2).

### LINCS CP benchmarking results (ensemble signatures)

We fit ensemble iPAS signatures for each multi-task value *λ*_2_ ∈ {0, 1, 2, 5, 10, 50, 100} using machine learning as described in the Methods. To benchmark their performance we followed a similar strategy as outlined above. For ensemble signatures, the evidence of pathway perturbation for a CP was defined as the linear combination of correlations with the iPAS signature for each cell line, where the weights of the combination are those found by machine learning. Again, we use AUROC as a summary of performance and summarize the results in Figure 3. Since we use signatures from each cell line, here we benchmarked using CPs in all cell lines. Notably, this is a tougher benchmarking than that of Figure 2 (where we used CPs in only the corresponding cell line) since this requires predicting pathway perturbation across a broad range of contexts.

**Figure 3:**
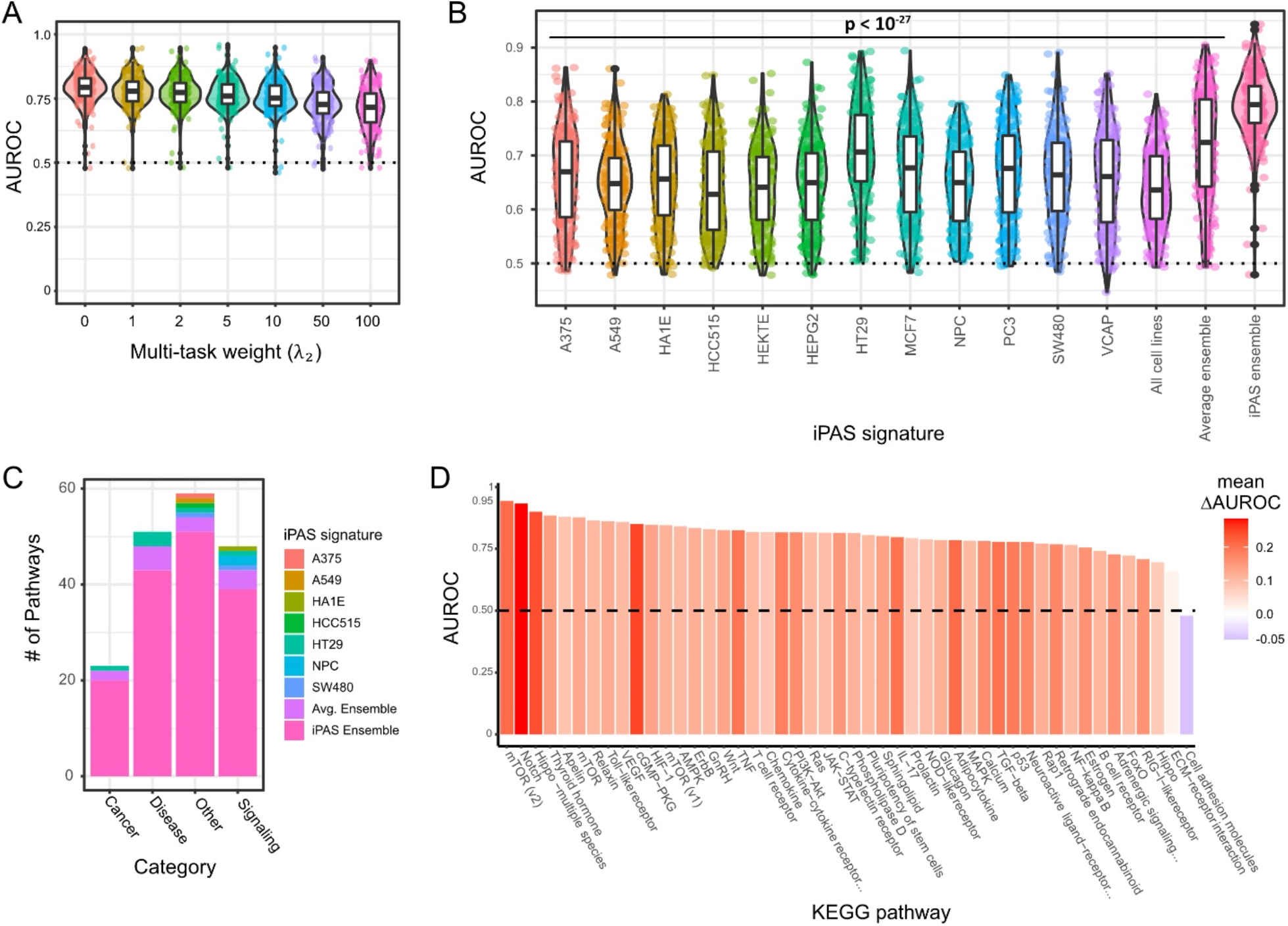
LINCS CP Benchmarking results for ensemble and individual iPAS signatures. A) Violin and box plots of AUROC for 181 KEGG pathways are shown for iPAS ensemble signatures created from multi-task signatures for each multi-task weight *λ*_2_ ∈ {1, 2, 5, 10, 50, 100}. B) Violin and box plots of AUROC for 181 KEGG pathways are shown for iPAS signatures (*λ*_2_ = 0) constructed using data from each cell line individually (A375, …, VCAP), using data from all cell lines together (“All cell lines”), using ensemble signatures using equal weights for each cell line (“Average ensemble”), and using ensemble signatures with optimal weights trained on LINCS CP data* (“iPAS ensemble”). *Optimal weights were trained using 2/3 of the LINCS CP data as a training set and ROC curves were constructed using the 1/3 of the data held out from training. C) A bar chart showing the number of pathways where each method performed best. We categorize KEGG pathways as either “Cancer”, “Disease”, “Signaling”, or “Other”. D) A bar chart showing AUROC values for all “Signaling” pathways. The color of the bars shows the average improvement in AUROC from pathway signatures trained on data from individual cell lines (A375, A549, …, VCAP) to the iPAS ensemble signatures.

We found that higher multi-task weights resulted in worse AUROCs for consensus signatures (Fig. 3A). This is because the ensemble learning method takes advantage of the diversity in data from different cell lines. As the multi-task term increases, the signatures are pushed closer to an average pathway signature and thus gradually lose their diversity. Since the ensemble signatures performed best for *λ*_2_ = 0, we compared these (labeled “iPAS ensemble”) to iPAS signatures from individual cell lines (*λ*_2_ = 0, “A375”, …, “VCAP”). We saw that the consensus signatures outperformed the individual ones by a large margin. Median area under the ROC curve (AUROC) for all 181 pathways was around 0.79 for consensus signatures and between 0.63 and 0.71 for individual cell lines (Fig. 3B).

We also tested two ways of naively combining data from the cell lines. First, we tested simply averaging the pathway consistency scores across all cell lines and using this as the pathway signature. This method performed similarly, but slightly worse than most of the individual signatures (“All cell lines”, median AUC around 0.63). Next, we tested naïve ensembles (“Average ensemble”) where we conserved the signatures from each of the 12 cell lines and instead averaged the *similarity* of a query signature with each of them. We found that the naïve ensembles performed slightly better than the iPAS constructed from individual cell lines, with a median AUROC of 0.72. However, the ensemble with trained coefficients (“iPAS ensemble”) significantly outperformed all of the iPAS from individual cell lines as well as the naïve ensembles (p < 10^−27^ for all individual comparisons, one-sided paired Wilcoxon rank-sum test).

We broke down results by individual pathways and found that the consensus signatures (“iPAS”) was the best performing signature method in the vast majority of pathways (Fig. 3C). We summarized these results for four categories of pathway (“Cancer”, “Disease”, “Signaling”, and “Other”) and found similar results for each. For example, in 39 out of 48 signaling pathways, the iPAS signatures performed best (Fig 3C).

In Figure 3D we show the AUROC results for iPAS ensemble signatures for each of the signaling pathways. For the mTOR pathway, we tested multiple variants: the original KEGG pathway (mTOR), a cleaned-up version of the pathway, with extra protein-protein interactions that were shown visually but missing from the underlying KEGG metadata (mTORv1), and our own curated version of the pathway (mTORv2). Among the best performing pathways are the mTOR (v2), Notch, and Hippo (multiple species) pathways, all of which saw a large improvement in AUROC (between 0.2 and 0.3) from the average individual iPAS to the ensemble iPAS.

### Benchmarking of mTOR pathway signatures

We show the ROC curves from the mTOR (v2) pathway (Fig 4A) for iPAS signatures from each method (all 12 individual cell lines, the two naïve combined methods, and the iPAS ensemble). The iPAS ensemble (light orange line) for the pathway has the highest AUC (0.94), but also demonstrates a very high sensitivity (y-axis) at high specificities (x-axis). This is the most crucial part of the ROC curve because the parts of the curve with low specificity relate to cutoffs that result in a large number of false positives. We also show precision-recall curves for the mTOR (v2) pathway (Fig 4B). Here, the contrast between the iPAS ensemble and the other methods is even more stark. The area under the curve (AUPR) is 0.479 for the iPAS ensemble and 0.105 for the second-best curve, the MCF7 iPAS. The performance of the average ensemble is much poorer here, coming in 6^th^ place with an AUPR of 0.026.

**Figure 4:**
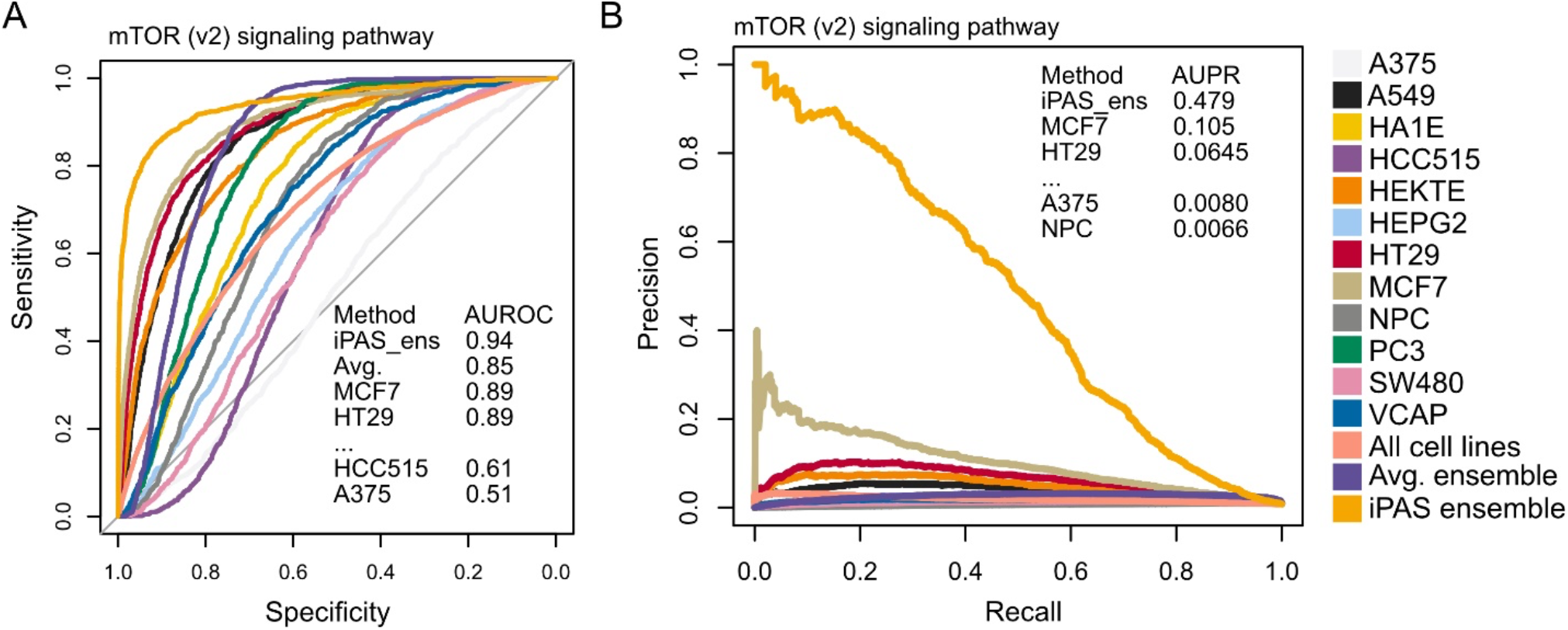
ROC and precision-recall plots for mTOR (v2) signaling pathway. A) ROC curves for the mTOR (v2) pathway for each method with selected areas under the curve (AUROC). B) Precision-recall curves for the mTOR (v2) pathway for each method with selected areas under the curve (AUPR).

### R package and web application for PAS analysis

iPAS is implemented as an open-source R package. The R package computes ensemble pathway scores for input signatures and computes z-scores and significance values via a permutation method. The package offers multiple methods for calculating pathway scores including Pearson correlation, cosine similarity, and the dot product. It also breaks down the scores in terms of the contribution of individual genes, so that the specific genes involved in pathway response can be identified. The package also faciliates visualization of results including heatmaps, bar charts, and plots showing the observed score vs. the null distribution of permutation scores..

## Discussion

The study of pathways in cancer is crucial to the success of the next generation of targeted therapies, such as kinase inhibitors (Arora and Scholar, 2005; Roskoski, 2015), that seek to effectively treat cancer while avoiding the extreme toxicity of chemotherapy. This field had considerable early success with Gleevec (imatinib) (Druker et al., 2001; Longo, 2017), an Abl kinase inhibitor that revolutionized treatment of chronic myelogenous leukemia (CML) starting in 2001, but has since largely failed to replicate this success (Hoelder et al., 2012; Janku et al., 2018). Researchers are currently in need of reliable methods to determine and quantify altered signaling pathway activity in cells. With proteomic data remaining more costly and less available than transcriptomic data for the near future, prediction of pathway activity from altered gene expression is a desirable prospect.

Many common pathway methods directly or indirectly rely on the assumption that transcript expression can be mapped to signaling molecules as a measure of activity. While pathway mapping methods have achieved success in some cases, their results overall have been mixed (Geistlinger et al., 2016). Recent efforts (Schubert et al., 2018; Ren et al., 2020) have shown that signature methods, derived from large expression databases of perturbation experiments, can outperform these methods in terms of pathway activity prediction. The LINCS L1000 dataset offers a significant resource of genetic and chemical perturbation response signatures from which we can identify genes whose expression is consistently altered upon pathway activation or inhibition.

In previous work (Ren et al., 2020) found that genetic knockdown signatures of pathway signaling molecules from LINCS could be used to create pathway signatures in each of twelve cell lines that could accurately predict chemical perturbation of pathways. In the tests, their methods outperformed the standard pathway mapping methods including SPIA (Tarca et al., 2009), CePa (Gu and Wang, 2013), and PathNet (Dutta et al., 2012).We developed the iPAS methodology in order to create more accurate pathway signatures by integrating data from many cell lines at once.

iPAS uses two strategies for data integration, unsupervised multi-task learning and supervised ensemble learning methods. Using the multi-task method, we created signatures that are specific to a single cell line but borrow genetic perturbation response information from all of the cell lines available, using a weight to determine the amount of influence from each of the two sources. Using our ensemble method we learned optimal consensus signatures, weighting the influence of the multi-task signatures pertaining to each cell line. In our performance benchmarking we found that both methods increased our accuracy in predicting pathway-targeting LINCS chemical perturbations over methods that used each cell line separately.

Overall, our findings show that iPAS is able to recover downstream transcriptional response genes from perturbation data across many cell lines with diverse tissues of origin and genomic alterations. By integrating data across cell lines, we find that we are able to improve our ability to predict signaling pathway alteration overall. Our methods improve on pathway mapping methods, which can only detect altered signaling when it is reflected in the expression of pathway members themselves, and not when it is only apparent in downstream transcriptional response genes. We also demonstrate the connection between genetic and chemical pathway perturbations.

The issue of whether to use cell line-specific signatures or consensus signatures requires careful consideration. Our results from ROC curves (Fig. 3A) clearly show that iPAS ensemble signatures are superior overall at identifying pathway-targeting chemical perturbations across a panel of twelve cell lines. The difference is particularly pronounced in some pathways when looking at precision-recall curves, for example in the mTOR pathway (Fig. 4B). However, one must remember that cell lines are separate model systems with distinct characteristics. Pathway response can be context-specific, so a signature from the same cell line or the same tissue may most accurately represent pathway activity in a certain context.In the context of exploratory research of large datasets and prioritizing chemicals for future research, ensemble signatures may be very useful due to their high overall performance in identifying pathway-targeting perturbations. Furthermore, their high performance over a panel of diverse cell lines suggests that they may be applicable to a wide array of additional cell lines outside of those used in this study, including many tissue contexts. At the same time, we need to keep in mind that the multi-task methodology is unsupervised, whereas ensemble method is supervised and may be less generalizable. The ensemble model may be over-fitted for the drugs currently profiled and for the drugs with many signatures in the data. On the other hand, the multi-task methodology never uses information from drug signatures and is completely based on the signatures of genetic perturbations of pathway genes. Therefore, it may generalize better to chemical perturbagens not profiled by LINCS.

We have shown that iPAS signatures can accurately predict pathway perturbation in a broad range of cellular contexts (breast, colon, kidney, liver, lung, melanoma, naso-pharyngial, and prostate). Our method combines techniques that exploit the similarity (multi-task learning) as well as the heterogeneity (ensemble learning) of pathway response among cell lines originating from diverse tissue contexts. Our results indicate that we improve our detection accuracy over past attempts as well as competing methods. We make our signatures easily available via an R package and web application. Looking forward, our methodology remains applicable to new high-resolution sequencing datasets, opening the possibility to even higher quality pathway signatures in the future.

## Acknowledgements

We would like to thank Dr. Jarek Meller for his insightful comments and suggestions on our methods.

