## Supplemental material for "Integrative signatures of signaling pathway response increase robustness and accuracy of pathway predictions"

### Supplementary methods

#### Processing of KEGG pathways

We use KEGG [1] pathway data from Ren et al. [2] as the basis for our pathways. To summarize, 328 homo sapiens KEGG pathways were downloaded using the KEGGRest R package. We keep pathways containing at least two "activation" or "inhibition" interactions and without conflicting interactions. For each pathway, a signed adjacency matrix was constructed. The adjacency matrix is defined as:

$$A_{ij} = \begin{cases} 1 \; if \; i \; and \; j \; share \; an \; "activating" \; edge \\ -1 \; if \; i \; and \; j \; share \; an \; "inhibiting" \; edge \\ 0 \quad if \; i \; and \; j \; do \; not \; share \; an \; edge \end{cases}$$

We group the pathways into categories as follows: a) pathways containing the word "cancer" are categorized as "cancer"; b) pathways related to another disease are categorized as "disease"; c) pathways containing the word "signal" or "signaling" are categorized as "signaling"; d) all other pathways are categorized as "other".

#### LINCS L1000 data

We summarize the LINCS data for chemical perturbations (CP) genetic perturbations (GP) in Figure S1. Out of all of the LINCS data[3-5], the chemical perturbations ("trt_cp") were by far the most numerous with over 300,000 signatures (Fig. S1A). There are also ~150,000 signatures of individual shRNAs ("trt_sh"). For constructing pathways signatures we used the "Consensus Gene Signatures" (CGS, "trt_sh.cgs), of which there were closer to 30,000 in total, over all cell lines. These "CGS" are consensus signatures created from 3 or more L1000 profiles of shRNAs targeting the same gene, but with different seed sequences. These are more reliable because large off-target effects corresponding to seed sequences were found in LINCS shRNA data[6]. Other types of LINCS signatures, over-expression (“trt_oe”), ligand (“trt_lig”), and CRISPR loss-of-function (“trt_xpr”) contained fewer signatures, making them less useful for our large-scale analyses.

Figure S1B breaks down the number of genetic perturbations (CGS, referred to GP in the main paper) per cell line. All of the data comes from custom microarrays called L1000, that measure the expression of 978 genes with bead-based probes. These ~1000 genes were chosen in a data-driven manner to be a minimal gene set that captures 80% of the information present in the transcriptome. The transcriptional data from L1000 was shown to be consistent with Affymetrix microarrays and RNA-sequencing[3]. The twelve LINCS cell lines that we use to make signatures are A375 (melanoma), A549 (lung), HA1E (kidney), HCC515 (lung), HEKTE (kidney), HEPG2 (liver), HT29 (colon), MCF7 (breast), NPC (naso-pharyngial), PC3 (prostate), SW480 (colon), and VCAP (prostate). Most cell lines contain between three and four thousand GPs. PC3 and MCF7 contain the most GP signatures, about five thousand, and NPC, HEKTE, and SW480 contain the least with less than a thousand each.

#### Modeling pathways using signed graphs

We model each pathway as a signed graph. In general, a signed graph $G = (V, W)$, consists of a set of vertices $V$ numbered from 1 to $m$ and a symmetric matrix $W$ (a weighted adjacency matrix) with zero diagonal entries. The non-zero entries in W represent a set of edges $E$ (pairs of vertices $(i, j)$ where $i, j \in V$) with weights $w_{ij} = w_{ji}$. We consider the edges of the graph to be un-directed, i.e. $(i, j) \in E$ implies $(j, i) \in E$, but signed so that inhibiting edges have a weight of -1 and activating edges have a weight of 1.

Since we consider only weights of 1, -1, or 0, the adjacency matrix for our graph is signed, but un-weighted: we denote it by $A$. Given the signed adjacency matrix, we define the signed Laplacian[7, 8] of the graph as $L = D - A$ where $A$ is defined as above and $D$ is a diagonal matrix with $D_{ii}$ equal to the number of edges connecting to node $i$. This Laplacian matrix can be thought of as a summary of the topology of the graph. It has a number of useful qualities that we can take advantage of.

#### Properties of the Laplacian matrix

The properties of un-directed, un-signed Laplacians and their connection to graph kernels and regularization are discussed in Smola and Kondor[9]. The properties are extended to signed Laplacians in Kunegis et al.[7] and Gallier[8]. The signed Laplacian is symmetric and positive semi-definite meaning that $x^T Lx \geq 0$ for all $x \in \mathbb{R}^m$, or equivalently, that all of the eigenvalues of $L$ are real and non-negative.

We consider vectors of expression of one gene at a time across pathway knockdowns. The quadratic form $x^T Lx = \sum_i \sum_j L_{ij}\, x_i x_j$ for an expression vector $x$ can be used as a measure of the "consistency" of $x$ with the pathway. A highly consistent expression vector $x$ will produce a value of $x^T Lx$ close to zero while a highly inconsistent vector will produce a large positive value. If we express the quadratic from in terms of the adjacency matrix $A$, this becomes more apparent.

$$x^T Lx = \frac{1}{2}\sum_i \sum_j |A_{ij}|\left(x_i - sgn(A_{ij}) \cdot x_j\right)^2 = \sum_{(i\sim j)^+} (x_i - x_j)^2 + \sum_{(i\sim j)^-} (x_i + x_j)^2$$

where $(i\sim j)^+$ means nodes $i$ and $j$ are connected by an activating edge and $(i\sim j)^-$ means nodes $i$ and $j$ are connected by an inhibiting edge.

We see that the quadratic form is a sum of squares for all interacting edges $e_{ij}$ connecting nodes $i$ and $j$. When $A_{ij}$ is an activating relationship, the summand is $(x_i - x_j)^2$, so *similar* expression values in a gene at these two nodes (knockdowns of interacting proteins) will produce a small or "consistent" value. When $A_{ij}$ is an inhibiting relationship, the summand is $(x_i + x_j)^2$, so *opposite* (up- vs. down-regulated) will produce a small or "consistent" value.

Furthermore, the eigenvectors of $L$ can give us useful information about the pathway. Since $L$ is positive semidefinite, its eigenvalues are all greater than or equal to zero: $0 \leq \lambda_1 \leq \lambda_2 \leq \dots \leq \lambda_m$. For a signed graph, the Laplacian has exactly one eigenvalue equal to zero if and only if it is connected and balanced [8]. The eigenvector corresponding to the eigenvalue of zero consists completely of 1's and -1's. This vector splits the graph into the two disjoint groups of nodes that can be thought of as "activators" and "inhibitors" of the pathway.

#### Pathway consistency scores

The first eigenvector (associated with an eigenvalue of zero) is the most "consistent" vector possible for the pathway: $v_1^T L v_1 = 0 \leq x^T L x$ for all $x \in \mathbb{R}^m$. Therefore, when training base signatures, we use only the first eigenvector of $L$ to score genes based on pathway consistency. We define the pathway consistency score as $S(x) = v_1^T x$ where $v_1$ is the first eigenvector of $L$ and $x$ is a vector of expression of one measured gene across knockdowns (GPs) in a pathway. For our base pathway signatures in each cell line, we score the L1000 genes by consistency of their expression with the pathway topology and select the best 100 genes, using their scores as a signature of pathway activity.

#### Pathway scoring for query signatures

We share the individual PAS signatures and ensemble PAS signatures in an R package (iPAS) along with some functions to compute pathway scores and significances. To score a query signature with a single PAS (e.g. from a specific cell line) we measure its similarity to the pathway signature. For a similarity, we use the Pearson correlation with each pathway signature:

$$r_{xy} = \frac{1}{n-1}\sum_{i=1}^{n}\left(\frac{x_i - \bar{x}}{s_x}\right)\left(\frac{y_i - \bar{y}}{s_y}\right)$$

where $x$ is the query signature, $y$ is the PAS, and $s_x = \sqrt{\frac{1}{n-1}\sum_{i=1}^{n}(x_i - \bar{x})^2}$ is the standard deviation of $x$ (defined similarly for $y$).

To score a pathway with an ensemble PAS, we use a linear combination of its scores with each individual PAS, where weights are equal to the ensemble weights that we derived using machine learning. So, for a query signature $x$, we let its similarity with the PAS in the $k$-th cell line be denoted $r_k = corr(x, y_k)$. Then the ensemble score is given by the linear combination

$$r = \sum_{k=1}^{N} \beta_k r_k = \beta_1 r_1 + \cdots + \beta_N r_N$$

To derive significance for these scores, we use a permutation method. We permute the query signature $n = 10{,}000$ times and calculate a pathway score for each permuted signature. We define a one-sided empirical p-value as $p = \frac{m}{n}$ where $m$ is the number of permuted scores more extreme than the observed score. We also define a pathway z-score as $z = \frac{s_{obs} - mean(s_{null})}{sd(s_{null})}$ where $s_{obs}$ is the observed pathway score and $mean(s_{null})$ and $sd(s_{null})$ are the mean and standard deviation of the null distribution (permutation) scores.

#### Gene importance scores

We can see that the Pearson correlation is equal to a constant times the dot product of the standardized (centered and scaled) versions of the query signature and the PAS.

$$r_{xy} = \frac{1}{n-1}\sum_{i=1}^{n} z_{xi} z_{yi} = \frac{1}{n-1} z_x^T z_y$$

This leads directly to a formula for the contribution of each gene to the pathway score in a single cell line:

$$c_i = z_{xi} z_{yi}$$

where $z_{xi}$ and $z_{yi}$ are, respectively, the values of the $i$-th gene in the query signature and the PAS after centering and scaling each.

To calculate the contribution of a single gene to an ensemble pathway score, we can simply combine the contributions to the score in each individual cell line, weighting each by the appropriate ensemble weight. If $c_{ik}$ is the contribution of gene $i$ to the pathway score in the $k$-th cell line, then their overall contribution to the ensemble pathway score is given by

$$c_i = \sum_{k=1}^{N} \beta_k c_{ik} = \beta_1 c_{i1} + \cdots + \beta_N c_{iN}$$

#### Multi-task signature methods

##### Overview of multi-task PAS signature creation

In addition to our base signatures and ensemble signatures, we try a method inspired by multi-task learning in order to "share" information between cell lines when creating pathway signatures. The idea is that pathway response is highly conserved and should be similar, but not exactly the same, in many cell lines. Rather than simply average gene expression over cell lines, we use a weight to push the expression toward a common average.

##### Background of multi-task learning

Multi-task learning is usually done in the setting of linear or logistic regression. Whereas single-task regression solves for one optimal vector of parameters (one "task"), multi-task regression jointly solves for separate optimal vectors for each of $N$ tasks. The data for single-task logistic regression consists of $n$ data points $\{(x_i, y_i)\}$ where $y_i \in \mathbb{R}$ is a response value and $x_i \in \mathbb{R}^P$ is an input value. In single-task linear regression, the objective is to find a vector of parameters that minimizes the following loss function:

$$\widehat{w} = \underset{w}{\arg\min} \sum_{i=1}^{n} (y_i - w^T x_i)^2 = \underset{w}{\arg\min} \|y - w^T X\|_2^2$$

It is common to add “regularization” terms to the loss function that limit the size of $w$. The most common of these are the $L_1$ or LASSO penalty, $\|w\|_1 = \sum_i |w_i|$, and the $L_2$ or ridge regression penalty, $\|w\|_2^2 = \sum_i w_i^2$. The $L_2$ penalty shrinks the values of $w$ toward zero while the $L_1$ penalty encourages sparsity in $w$. Combined with weights $\lambda_1$ and $\lambda_2$, these penalties form what is called Elastic Net regression:

$$\widehat{w} = \underset{w}{\arg\min} \|y - w^T X\|_2^2 + \lambda_1 \|w\|_1 + \lambda_2 \|w\|_2^2$$

In the multi-task setting, we may have data from $t = 1, \dots, N$ separate tasks with $i = 1, \dots, n_t$ data points each $\{(x_{it}, y_{it})\}$:

$$\widehat{W} = [w_1 \quad \cdots \quad w_N] = \underset{\{w_1, \ \dots, \ w_N\}}{\arg\min} \sum_{t=1}^{N} \sum_{i=1}^{n_t} (y_{it} - w_t^T x_{it})^2 = \underset{\{w_1, \ \dots, \ w_N\}}{\arg\min} \sum_{t=1}^{N} \|y_t - w_t^T X_t\|_2^2$$

Often there are regularization terms added onto the end of this loss function:

$$\widehat{W} = [w_1 \quad \cdots \quad w_N] = \underset{\{w_1, \ \dots, \ w_N\}}{\arg\min} \sum_{t=1}^{N} \|y_t - w_t^T X_t\|_2^2 + \lambda_1 \Omega(W) + \lambda_2 \|W\|_F^2$$

The first penalty, $\Omega(W)$, is a penalty that encourages similar solutions among the $w_t$, which can take many forms. The second penalty, is the square of the Frobenius norm of the matrix $W$, which is simply the sum of squares of all of its entries, $\|W\|_F^2 = \sum_i \sum_j w_{ij}^2$. This is just the sum of the $L_2$ penalties for each vector, which shrinks their values toward zero as above.

#### Multi-task signature methods

For these signatures, we set up a minimization problem with a weight that pushes expression in individual cell lines toward an overall average, similar to a multi-task learning problem where the regularization term encourages similar solution vectors. For a given pathway with GP signatures of $P$ pathway nodes, we consider the expression of each gene separately. We let $y_{ik}$ be the vector of expression of gene $i$ in the $k$-th cell line across all $P$ pathway nodes, $y_{ik}^T = [y_{i1k} \quad \cdots \quad y_{iPk}]$. We look at its expression across all $N$ cell lines: $y_i = vec(Y_i)$ where $Y_i = [y_{i1} \quad \cdots \quad y_{iN}]$. $Y_i$ is a matrix where each column is the expression of gene $i$ across pathway nodes in one cell line and $y_i$ is the "vectorized" version of this matrix, a vector in which the columns of $Y_i$ are stacked on top of each other.

We want to pull each target vector $y'_{ik}$ towards a common mean $\bar{y}'_i = \frac{1}{N}\sum_{k=1}^{N} y'_{ik}$, encouraging similar solutions in each cell line.The minimization problem is set up as follows:

$$\hat{y}_i = \begin{bmatrix} \hat{y}_{i1} \\ \vdots \\ \hat{y}_{iN} \end{bmatrix} = \underset{\{y'_{i1}, \ \ldots, \ y'_{iN}\}}{\arg\min} \sum_{k=1}^{N} \|y_{ik} - y'_{ik}\|_2^2 + \lambda_1 \sum_{k=1}^{N} {y'_{ik}}^T L y'_{ik} + \lambda_2 \sum_{k=1}^{N} \|\bar{y}'_i - y'_{ik}\|_2^2$$

where $y'_{ik}$ is a target vector in cell line $k$ and $\{\hat{y}_{i1}, \ldots, \hat{y}_{iN}\}$ is the solution that minimized the right-hand side.

To find this solution to this problem, we combine all of the cell lines into one vector and write each term as matrix-vector multiplication. Similarly to $y_i$ defined above, let $y'_i = vec(Y'_i)$ where $Y'_i$ is the matrix $Y'_i = [y'_{i1} \quad \ldots \quad y'_{iN}]$. Also, let $\bar{\bar{y}}' = vec([\bar{y}'_i \quad \cdots \quad \bar{y}'_i])$ and $\tilde{L} = \begin{bmatrix} L & \cdots & 0 \\ \vdots & \ddots & \vdots \\ 0 & \cdots & L \end{bmatrix}$.

Then we can re-write the above equation without the sums as:

$$\hat{y}_i = \begin{bmatrix} \hat{y}_{i1} \\ \vdots \\ \hat{y}_{iN} \end{bmatrix} = \underset{\{y'_{i1}, \ \dots, \ y'_{iN}\}}{\arg\min} \ \|y_i - y'_i\|_2^2 + \lambda_1 {y'_i}^T \tilde{L} y'_i + \lambda_2 \left\|\overline{\overline{y'}} - y'_i\right\|_2^2$$

This has solution

$$\hat{y}_i = \left(I + \lambda_1 \tilde{L} + \lambda_2 (I - B)\right)^{-1} y_i$$

where $B$ is a matrix such that $By'_i = \overline{\overline{y'}}$.

Defined in this way, $B$ is an $NP \times NP$ matrix $B = \frac{1}{N} \begin{bmatrix} I_P & \cdots & I_P \\ \vdots & \ddots & \vdots \\ I_P & \cdots & I_P \end{bmatrix} \Bigg\} N \ times$

The first term keeps the target vector near the observed values while the second term encourages a vector that "agrees" with the pathway topology, and the third term pushes the target vector for each cell line toward an overall average. We can call $\hat{y}_{ik}$ the "regularized" expression of gene $i$, in cell line $k$, across pathway nodes for regularization weights $\lambda_1$ and $\lambda_2$. After jointly solving for $\hat{y}_{ik}$ for each cell line, we score this regularized expression by the same metric as before: $S(\hat{y}_{ik}) = v_1^T \hat{y}_{ik}$ where $v_1$ is the first principal component of the pathway Laplacian.

After creating scores for each gene, in each cell line, for the pathway we use these scores in the creation of our "signatures" of pathway activity. As with the base signatures, we use the scores of the top 100 genes as a signature for the pathway. We create signatures in this way for a range of regularization weights $\lambda_1, \lambda_2 \in \{0, 1, 2, 5, 10, 50, 100\}$. When $\lambda_2 = 0$, there is no sharing of information across cell lines and the signatures are essentially created separately. When $\lambda_2 = 100$, the expression vector for each cell line is pushed almost completely to the mean over all cell lines.

#### Ensemble signature methods

For logistic regression, the log-odds of a set of correlations belonging to the "Case" group vs. the "Control" group is modeled as a linear combination of the correlations in each cell line:

$$\log \frac{\Pr(G = Case|\, X = x)}{\Pr(G = Control \,|\, X = x)} = \beta_0 + \beta^T x = \beta_0 + \beta_1 x_1 + \cdots + \beta_m x_m$$

Where $\Pr(G = Case|\, X = x) = \frac{e^{\beta_0 + \beta^T x}}{1 + e^{\beta_0 + \beta^T x}}$ and $\Pr(G = Control|\, X = x) = \frac{1}{1 + e^{\beta_0 + \beta^T x}}$

For training, we use two-thirds of the LINCS CP signatures targeting the pathway, saving the other one-third for testing. We minimize the following objective functional, the negative binomial log-likelihood (where the response $y_i = 1$ for cases and $y_i = 0$ for controls), with respect to the parameters $\beta_0, \beta$ for a given LASSO penalty weight $\lambda$.

$$\min_{\beta_0, \beta} - \left[ \frac{1}{N} \sum_{i=1}^{N} y_i \cdot (\beta_0 + \beta^T x_i) - (1 - y_i) \cdot \log\left(1 + e^{(\beta_0 + \beta^T x_i)}\right) \right] + \lambda \sum_{j=1}^{m} |\beta_j|$$

We use the *glmnet* R package to solve this functional along a range of lambda values using 10-fold cross-validation. Using the AUC of an ROC curve as our measure of deviance, we select the largest value of lambda that is within one standard error of the minimum misclassification error. The output of the model is an optimal set of values for the coefficients $\beta_0$ and $\beta$, weighting the correlation of an input signature with the pathway's PAS from each cell line.

### Supplementary results

#### Benchmarking performance of multi-task PAS signatures

We benchmarked the multi-task PAS signatures in the same way as the individual and ensemble signatures, although here we benchmark using CPs from only the corresponding cell line. In the main text, we benchmark signatures against CPs from all 12 cell lines, which can be seen as a more challenging test of performance. Figures S2-S3 summarize the performance of multi-task signatures for different choices of $\lambda_1$ and $\lambda_2$ by the area under the ROC curves (AUROC). Varying the pathway topology parameter ($\lambda_1$) did not make a difference to the performance of signatures (Fig. S2). However, increasing the across multi-task (cell-line sharing) term ($\lambda_2$) from 0 to 100 did increase performance slightly in almost all of the cell lines (Fig. S3).

### Supplemental Figures

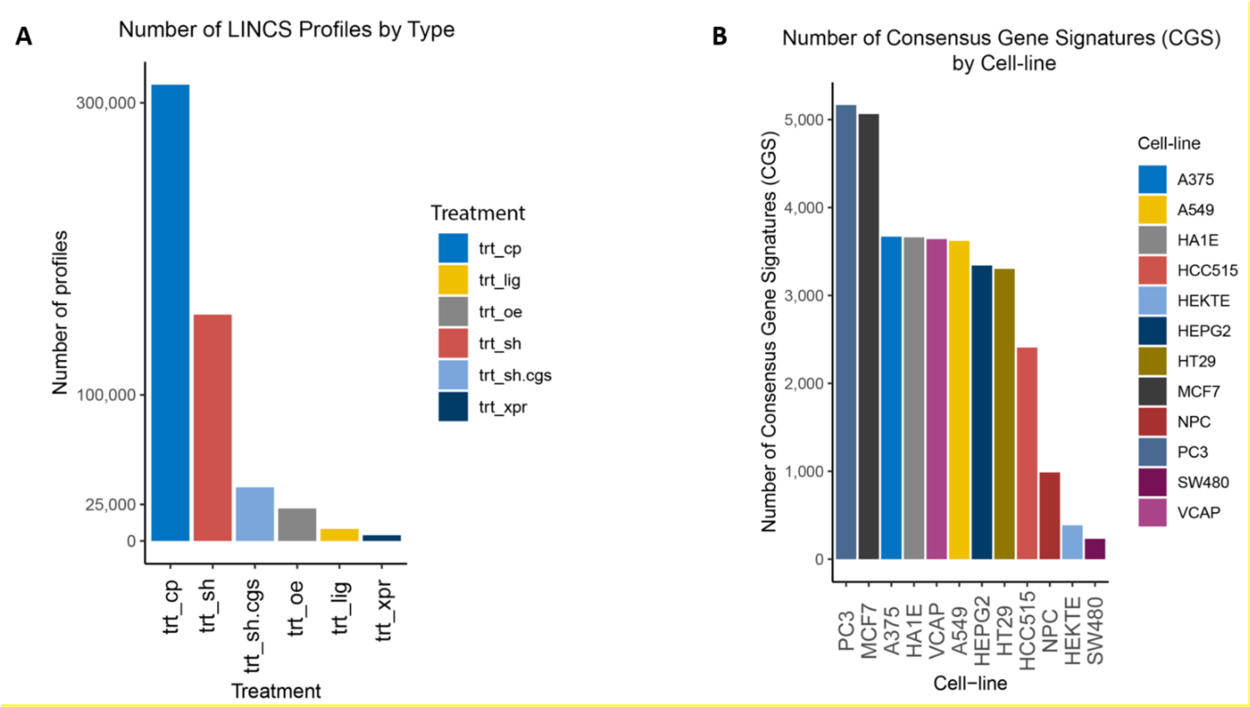


**Figure S1: Summary of LINCS data.** A) The number of LINCS signatures by type. trt_cp = Chemical perturbagen (Level 5), trt_sh = individual shRNA loss-of-function signatures (Level 4), trt_sh.cgs = replicate consensus shRNA signatures ("Consensus Gene Signatures - CGS", Level 5), trt_oe = over-expression signatures, trt_lig = ligand perturbation signatures, trt_xpr = CRISRP loss-of-function signatures. B) The number of LINCS shRNA consensus signatures (CGS) by cell line.

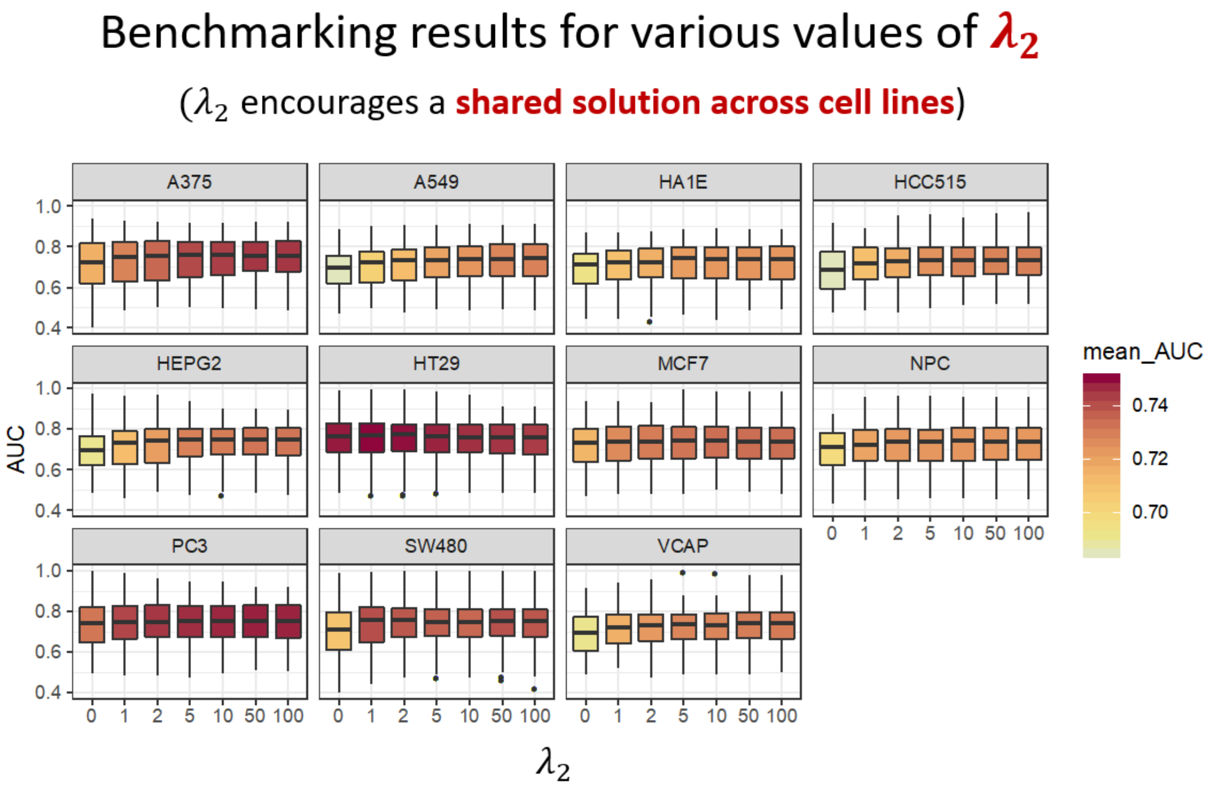


**Figure S2: Benchmarking results for joint/multi-task signatures using various cell-line multi-task weights.** The y-axis is the area under the curve (AUC) of ROC curves. Each boxplot represents AUCs for PAS signatures from 181 KEGG pathways. The color of the boxplot shows the average AUC for that group of points. The results are divided by cell line and the weight of the pathway topology term (ranging from 0 to 100) is shown along the x-axis.
